# FlyPhoneDB: An integrated web-based resource for cell-cell communication prediction in *Drosophila*

**DOI:** 10.1101/2021.06.14.448430

**Authors:** Yifang Liu, Yanhui Hu, Joshua Shing Shun Li, Jonathan Rodiger, Aram Comjean, Helen Attrill, Giulia Antonazzo, Nicholas H. Brown, Norbert Perrimon

## Abstract

Multicellular organisms rely on cell-cell communication to exchange information necessary for developmental processes and metabolic homeostasis. Cell-cell communication pathways can be inferred from transcriptomic datasets based on ligand-receptor (L-R) expression. Recently, data generated from single cell RNA sequencing (scRNA-seq) have enabled L-R interaction predictions at an unprecedented resolution. While computational methods are available to infer cell-cell communication in vertebrates such a tool does not yet exist for *Drosophila*. Here, we generated a high confidence list of L-R pairs for the major fly signaling pathways and developed FlyPhoneDB, a quantification algorithm that calculates interaction scores to predict L-R interactions between cells. At the FlyPhoneDB user interface, results are presented in a variety of tabular and graphical formats to facilitate biological interpretation. To demonstrate that FlyPhoneDB can effectively identify active ligands and receptors to uncover cell-cell communication events, we applied FlyPhoneDB to *Drosophila* scRNA-seq data sets from adult midgut, abdomen, and blood, and demonstrate that FlyPhoneDB can readily identify previously characterized cell-cell communication pathways. Altogether, FlyPhoneDB is an easy-to-use framework that can be used to predict cell-cell communication between cell types from scRNA-seq data in *Drosophila*.

## INTRODUCTION

Single cell RNA sequencing (scRNA-seq) is increasingly being used for high-throughput and high-precision studies to characterize cell states and cell types, and in particular is quickly becoming the method of choice to study developmental and physiological processes. ScRNA-seq avoids the limitation of only detecting average expression levels of RNAs as in traditional bulk RNA-seq approaches, and allows studies of sample heterogeneity at the single-cell level. ScRNA-seq methods have also improved in terms of throughput and scalability, such that it is now possible to analyze tens of thousands of cells in one experiment (Fan *et al*. 2015; Klein *et al*. 2015; Macosko *et al*. 2015). Further, improvements in throughput and accuracy have expanded potential applications of scRNA-seq technology. For example, scRNA-seq is gradually being used to gain a deep understanding of entire development processes, including, through cluster analysis of all cells, definition of new subgroups, comparisons of subgroup heterogeneity, reconstruction of pseudo-time trajectories, generation of regulatory networks during development processes, and prediction of cell-cell interactions (Ghosh *et al*. 2020; Hung *et al*. 2020; Tattikota *et al*. 2020).

Several methods have been developed to predict cell-cell communication from scRNA-seq data, including CellPhoneDB, NicheNet, and CellChat (Browaeys *et al*. 2020; Efremova *et al*. 2020; Jin *et al*. 2021). All of these methods start by generating a single-cell gene expression matrix of ligand-receptor (L-R) pairs that is then used to predict the strength of interactions between cells. Each method has its own strengths. CellPhoneDB in its predictions of cell-cell communication not only considers L-R interactions but also considers co-expression expression of components of multi-subunit L-R complexes (Efremova *et al*. 2020). NicheNet adds another layer of complexity as it predicts ligand-target interactions by integrating L-R information with downstream signal transduction and gene regulatory network information (Browaeys *et al*. 2020). CellChat considers signal cofactors, including inhibitory and stimulatory membrane-bound coreceptors, soluble agonists, and antagonists of cell-cell communication (Jin *et al*. 2021). All of these tools rely on the annotation of ligands and receptors and some rely on annotation of signaling pathways. Only CellChat supports analysis of *Drosophila* data but is not easy to use for this purpose. For example, preparing files in the format needed for analysis requires extra work and it does not allow users to view activities of individual core pathway components.

Here, we developed FlyPhoneDB, a quantification algorithm that calculates interaction scores to predict L-R interactions between cells in *Drosophila* (Figure 1a). We first established a high confidence list of L-R pairs specific for the major *Drosophila* signaling pathways and stored the annotation in FlyPhoneDB. Next, we developed a pipeline that calculates interaction scores in scRNAseq data, which is then used to make predictions of L-R interactions between cells. We demonstrate the utility of the tool by analyzing three published scRNAseq datasets from the adult *Drosophila* midgut, abdomen, and immune system (Ghosh *et al*. 2020; Hung *et al*. 2020; Tattikota *et al*. 2020). Finally, we developed a web-based FlyPhoneDB user interface (http://www.flyrnai.org/tools/fly_phone), where users can browse data, upload additional L-R pairs, and analyze their own data sets.

**Figure 1.**
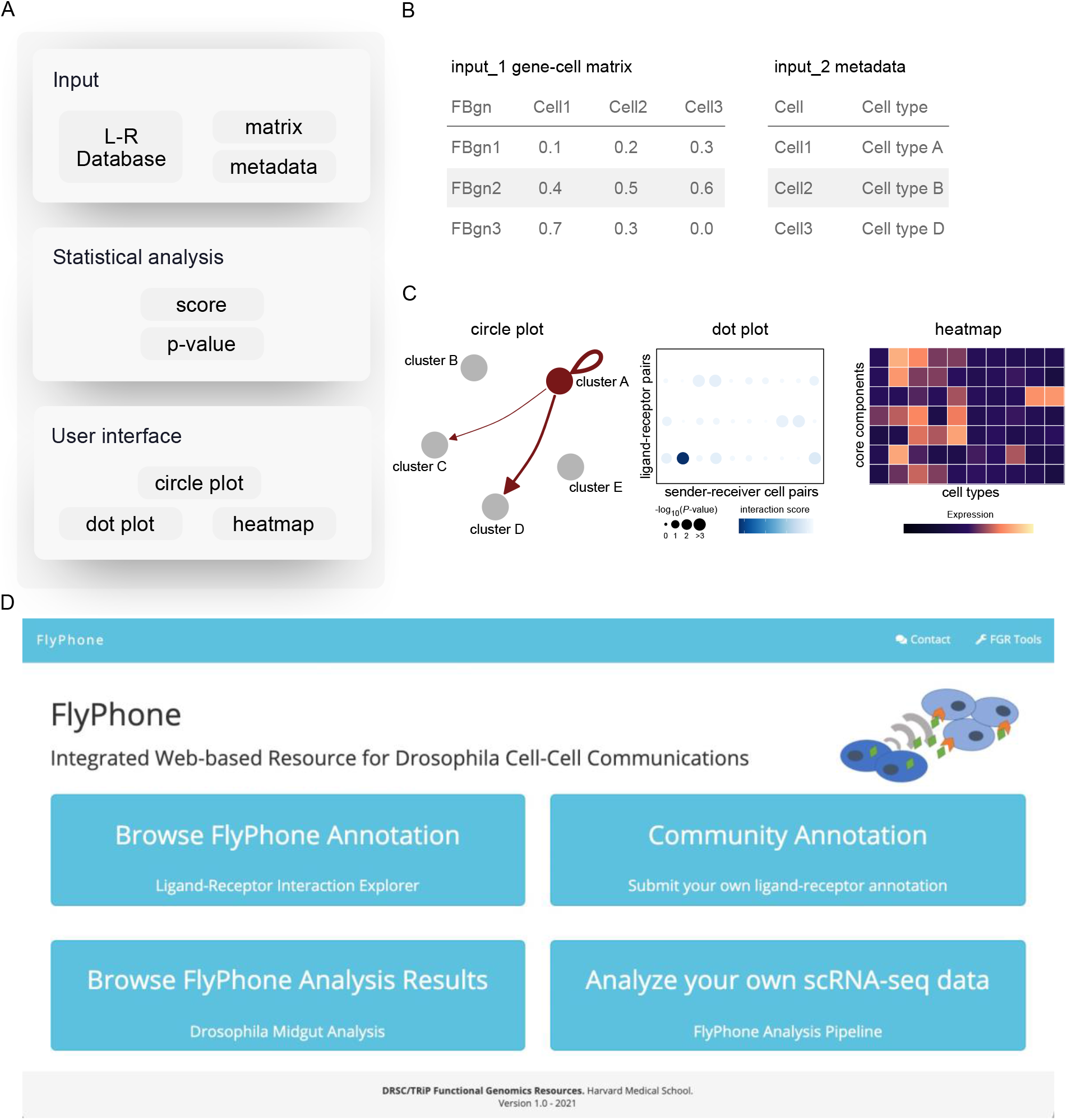
Overview of FlyPhoneDB. (A) Framework of FlyPhoneDB with input, statistical analysis, and user interface. (B) FlyPhoneDB requires users to provide gene-cell matrix and metadata that contains barcode and cluster information. (C) FlyPhoneDB provides various visualization tools to show ligand-receptor interactions between cell types. (D) FlyPhoneDB website contains four different sections: 1.) Browse the annotation database; 2.) Submit your own annotations; 3.) Browse an example; and 4.) Analyze new datasets.

## Materials and Methods

### Manual curation of ligand-receptor interactions for inclusion in FlyPhoneDB

To establish a high-confidence list of *Drosophila* L-R pairs, we manually curated information in the GLAD database (Hu *et al*. 2015; Upadhyay *et al*. 2017; Gontijo and Garelli 2018), FlyBase (Larkin *et al*. 2021) and QuickGO (Huntley *et al*. 2015) and imported these data into FlyPhoneDB.

Other core components of major signaling pathways, such as transcription factors, were directly imported from GLAD.

### Processing of Drosophila scRNA-seq data

The *Drosophila* midgut scRNA-seq dataset reported by Hung et al. (2020) was chosen for this study because a number of signaling pathways that maintain gut homeostasis have been well characterized, thus allowing us to evaluate the accuracy of FlyPhoneDB. The gene expression matrix and metadata were retrieved from GEO (accession code: GSE120537). The gene expression matrix was stored in the file “GSE120537_counts.csv.gz” in which rows are genes, columns are barcodes, and values are raw counts. The metadata was stored in the file “GSE120537_metadata.csv.gz” in which barcodes and cell type columns were extracted from this file (Figure 1b). This information can also be extracted from Seurat Object with the function GetAssayData and seuratObj@meta.data (Stuart *et al*. 2019). The abdomen and blood datasets were obtained from GEO (GSE147601 and GSE146596, respectively) and processed similarly. The wild type condition was analyzed from abdomen dataset and the wounded condition was analyzed from blood dataset.

### Calculation of L-R interaction scores and specificity

Expression values for each gene were normalized by dividing them by the total expression in each cell and then multiplying by the scale factor 10,000. Ligand and receptor expression levels were extracted from this normalized matrix based on the L-R pair database. Next, the average ligand and receptor expression values for each cell type were calculated by combining cell type information from the input metadata with the L-R expression matrix. The interaction score was calculated as the product of log transformed average ligand expression plus a pseudocount of 1 in the ‘sender cell’ with the log transformed average receptor expression plus a pseudocount of 1 in the ‘receiver cell’. Specificity was calculated using a permutation test by random shuffling of the original cell type assignments from the metadata (1,000 times by default) and then recalculating the interaction scores. P-values were computed based on the interaction score distribution of randomly shuffled cell types. P-values < 0.05 were considered significant.

### Construction of the database and website

The L-R pairs and core components information was stored in a MySQL database. The back end of the website was written in PHP and the front end was written in HTML. The JQuery JavaScript library and the DataTables plugin were used in user interface and displaying tables. Both the database and website are hosted on the O2 high-performance computing cluster at Harvard Medical School that is supported by Research Computing group.

#### Data Availability Statement

FlyPhoneDB web server is available at https://www.flyrnai.org/tools/fly_phone/web/. Three datasets used in this study are accessible through GEO series accession number GSE120537 (https://www.ncbi.nlm.nih.gov/geo/query/acc.cgi?acc=GSE120537), GSE146596 (https://www.ncbi.nlm.nih.gov/geo/query/acc.cgi?acc=GSE146596), GSE147601 (https://www.ncbi.nlm.nih.gov/geo/query/acc.cgi?acc=GSE147601).

### Code availability

The code is available upon request from the author.

## Results

### Overview of FlyPhoneDB

FlyPhoneDB uses single-cell gene expression matrix and metadata that contains cell annotation as the input to calculate interaction scores based on gene expression and L-R pairs. To implement FlyPhoneDB, we first manually curated signaling pathway information from GLAD, FlyBase and QuickGO to generate a list of high confidence L-R pairs. This resulted in a set of 196 L-R pairs with the majority representing the EGFR, PVR, FGFR, Hedgehog, Hippo, Insulin, Notch, JAK/STAT, TGF-β, TNFα, Wnt, Toll, and Torso signaling pathways. Next, to identify the specificity of L-R interactions, FlyPhoneDB permutes the cell annotation and recalculates the interaction score. Scores with P-values of < 0.05 were considered significant. The lower the P-value, the more specific the interaction between two cell clusters is predicted to be.

The FlyPhoneDB user interface supports a variety of visualizations, including circle plots, dot plots, and heatmaps (Figure 1c). Circle plots are provided for each signaling pathway. These depict all of the potential cell-cell communication events, with each node representing a unique cell type and each edge representing a communication event. The thickness of an edge reflects the interaction strength of the communication event, such that users can quickly identify cell types of interest for each pathway. Dot plots are designed to provide additional detail. For example, in cases where there are multiple ligands and/or receptors involved in a pathway we use one dot plot to illustrate the statistical outputs for all of the L-R pairs between any two cell types. After users identify pathways and related cell types from a circle plot, users can zoom into more detailed information on the relevant dot plots. Heatmaps, based on the average expression level of additional major components, i.e., other than ligands and receptors, in all cell types and for each pathway are also provided. Lists of additional major components for each pathway were obtained from the MIST database (Hu *et al*. 2018). These heatmaps provide a quick way for users to compare expression levels among different cell types. We also provide thumbnails of the main pathways in EGFR, FGFR, Insulin, Pvr, Torso, TNFα, TGF-β/BMP, TGF-β/Activin, Notch, Wnt-TCF, Hedgehog, JAK-STAT, Hippo, and Toll, which will be available at FlyBase (http://flybase.org/; release FB2021_04) (Figure 3).

### FlyPhoneDB website

The FlyPhoneDB site contains four different sections that allow users to do the following (Figure 1d); 1.) Browse the annotation of L-R pairs as well as the core component for each pathway with an option to download the information (Figure 2); 2.) Provide their own annotations by uploading a file with new information. Users have the option to upload one pair at a time or upload more than one pair in batch mode using a template file. An upload triggers an email message to FlyPhoneDB. The information will then be reviewed by staff and updated on the backend database; 3.) Browse the analysis result of scRNA-seq dataset from the *Drosophila* midgut as user case. Users can browse L-R interactions illustrated by dot plots and view expression patterns of all core components on a heatmap; and 4.) Analyze new datasets. Users can upload an expression matrix file obtained from a scRNA-seq dataset and the corresponding metadata file, and then analyze the dataset using the FlyPhoneDB pipeline. An email message is sent automatically when the analysis is completed and result files, including a table of statistics and visualizations, is made available for download.

**Figure 2.**
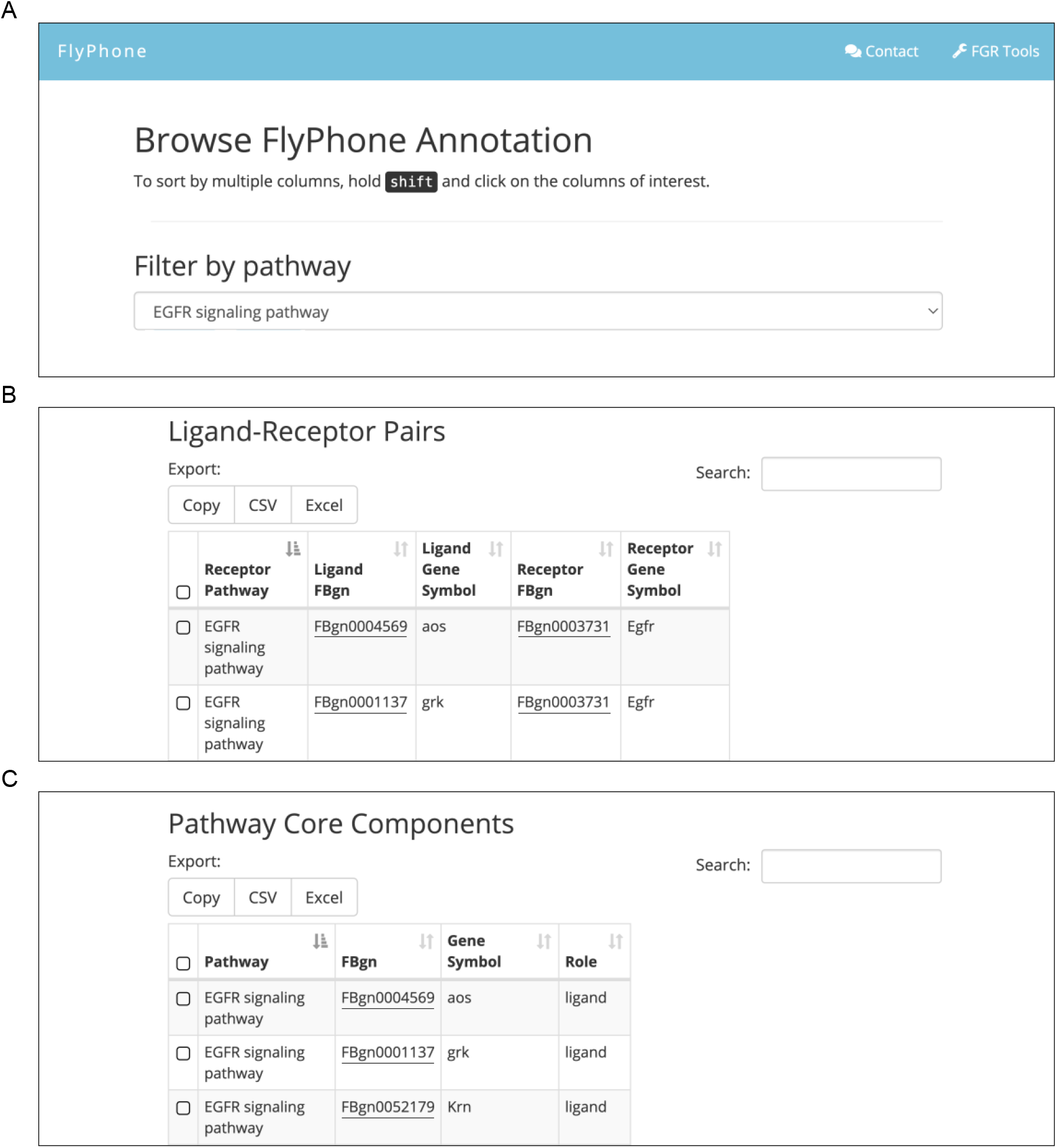
FlyPhoneDB annotation panel. (A) Browser of FlyPhone annotation. (B) Annotation of L-R pairs. (C) Core components information for each signaling pathway. Users can filter, sort, and download the various tables.

**Figure 3.**
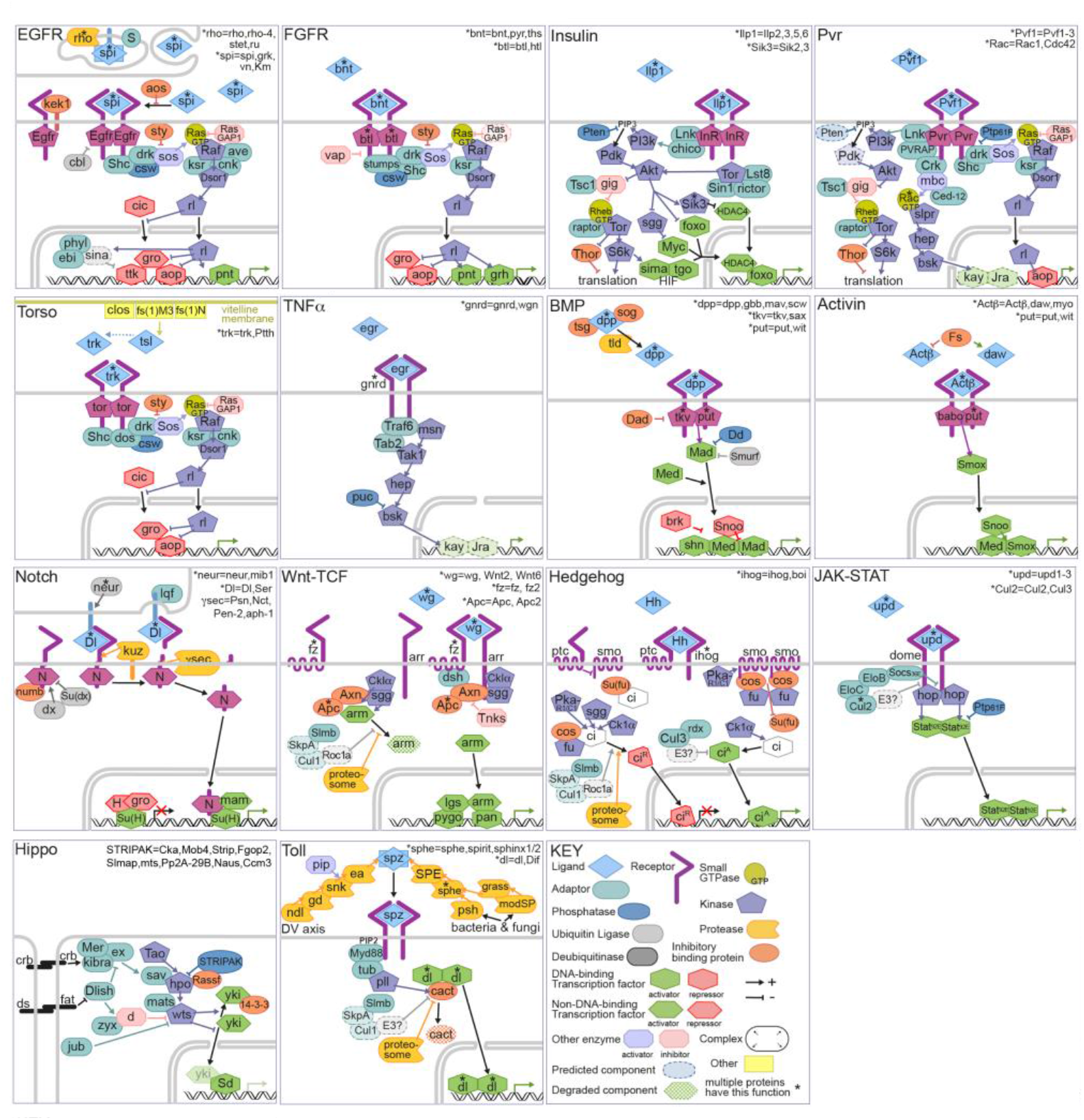
Thumbnails of the main signaling pathways. A set of thumbnails representing EGFR, FGFR, Insulin, Pvr, Torso, TNFα, TGF-β/BMP, TGF-β/Activin, Notch, Wnt-TCF, Hedgehog, JAK-STAT, Hippo, and Toll signaling pathways.

### Using FlyPhoneDB to identify signaling pathways in adult *Drosophila* organs

To assess FlyPhoneDB, we tested whether the tool could predict from existing scRNA-seq data sets cell-cell communication pathways that have been reported in the literature. We first focused our analysis on the fly gut, as a number of specific signaling pathways have been shown to regulate the maintenance of the adult midgut in an autocrine or paracrine manner.

The adult midgut largely consists of absorptive enterocytes (ECs) and secretory enteroendocrine cells (EEs) that are replenished by proliferative intestinal stem cells (ISCs). These three major cell types can be further categorized into sub-types according to their spatial or transient states (Buchon *et al*. 2013; Dutta *et al*. 2015; Guo *et al*. 2019; Hung *et al*. 2020; Hung *et al*. 2021). Previously, we identified 22 clusters in scRNA-seq data acquired from whole *Drosophila* midguts (Hung *et al*. 2020). These clusters included one cluster annotated as ISC/EBs, 14 clusters annotated as ECs, three clusters annotated as EEs, one cluster annotated as cardia, and three clusters of unidentified cell types. Among the EC clusters, four clusters (aEC1-4) correspond to the anterior midgut, one (mEC) maps to the middle midgut, three (pEC1-3) to the posterior, one to copper and iron cells, one to large flat cells (LFC), and one to differentiating EC (dEC), and three were annotated as ‘EC-like’ clusters. We applied FlyPhoneDB to this dataset and constructed heatmaps to visualize expression levels for each of the 13 major signaling pathways. We also generated visual representations of the interaction score and statistical significance of all possible combinations of L-R cell-cell interaction pairs.

We first looked at the Notch signaling pathway (Figure 4). ISCs produce the ligand *Delta* (*Dl*), which activates the Notch (N) receptor, triggering ISCs to develop into EBs (Micchelli and Perrimon 2006; Ohlstein and Spradling 2006). Subsequently, EBs differentiate towards the EC lineage. Accordingly, our expression profile shows enrichment in the ISC/EB cluster of *Dl, N*, and other key components of the Notch signaling pathway, including *kuzbanian* (*kuz*), an ADAM family metalloprotease that plays a role in the cleavage of N (Lieber *et al*. 2002), and many of the *enhancer of split* (*E(spl)*) genes. Additionally, ISC/EB>ISC/EB signaling had the strongest Dl-N interaction score of all pairwise interactions.

**Figure 4.**
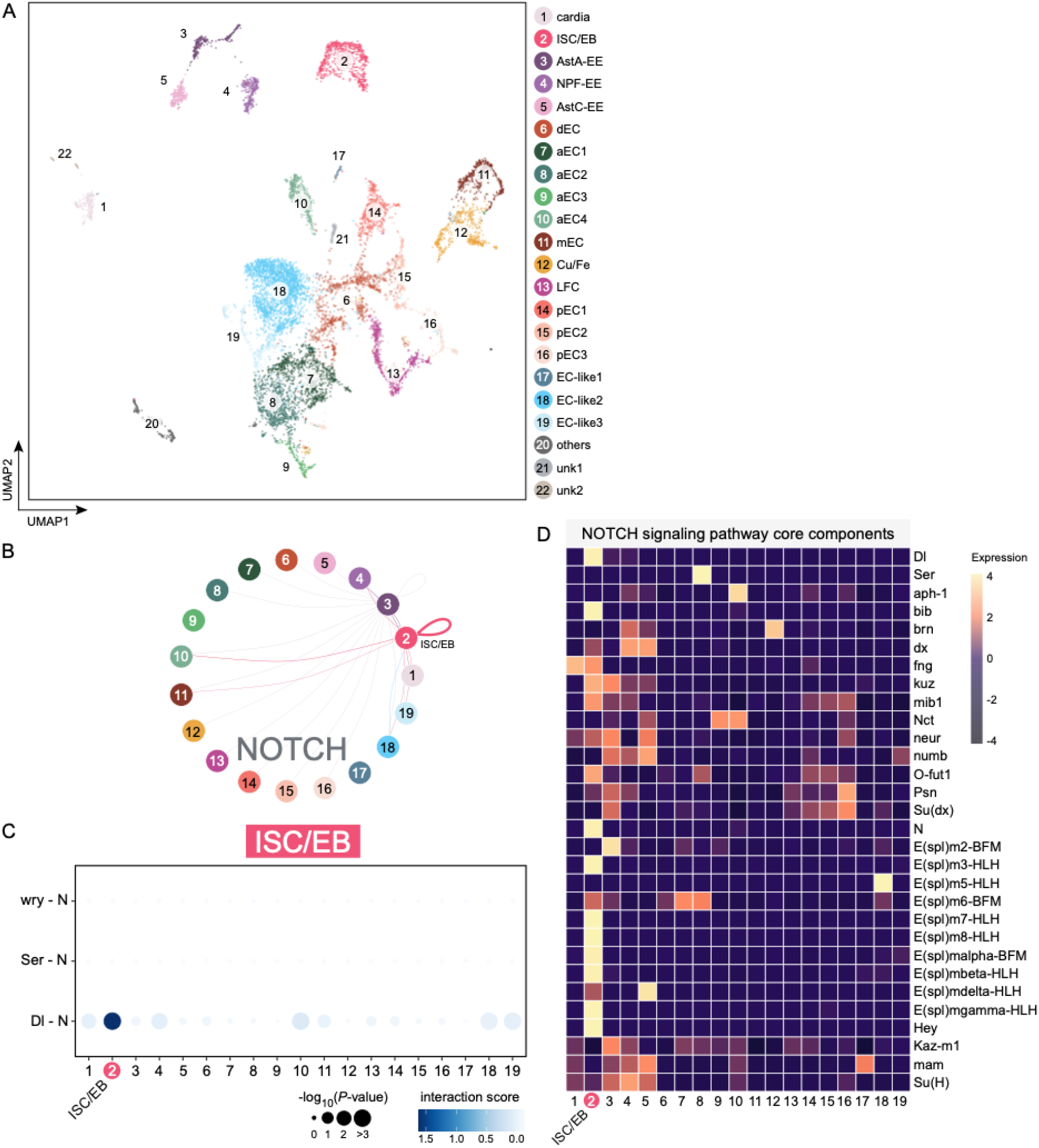
Analysis of signaling pathways in the adult *Drosophila* midgut. (A) UMAP of the *Drosophila* midgut (data from Hung *et al*. 2020). (B) Circle plot showing the significant interaction between the Delta (Dl) ligand and Notch (N) receptor in ISC/EB. (C) Dot plot showing the significant ligand-receptor pairs of Dl-N in ISC/EB. (D) Heatmap showing that Notch signaling in the entire gut is used only in ISC/EBs. It also gives a view of the components of the Notch network that is being used.

We next looked at the FlyPhoneDB output related to the JAK/STAT pathway. The L-R pairs for this pathway consist of binding interactions between any of the three Unpaired ligands (Upd1-3) and the Domeless (Dome) receptor. In the gut, all Upds are upregulated upon injury (Jiang *et al*. 2009). Specifically, Upds produced from ECs activate JAK/STAT signaling in Dome-expressing ISCs to promote cell division. Upds are also required for proliferation of ISCs under homeostatic conditions (Osman *et al*. 2012). Accordingly, we observed *upd2-3* expression in the aEC1, mEC, and pEC2-3 clusters, and *dome* expression in the ISC/EB cluster (Supplementary figure 1A). Notably, for the L-R pair Upd3-Dome, the cell interaction score for aEC1 and pEC3 cluster signaling to the ISC/EB cluster was greater than that of ISC/EB signaling to these EC clusters. This indicates that the directionality of signaling can be inferred from the cell interaction scores. We also found that ISC/EB to ISC/EB signaling had significant interaction scores for both the Upd2-Dome and Upd3-Dome L-R pairs, suggesting an autocrine role for Upd2 and Upd3.

We next looked at the EGFR signaling pathway. Upon damage or infection of the midgut, EGFR signaling in ISCs is enhanced by the upregulation and secretion of the EGFR ligands Vein (Vn), Spitz (Spi) and Kernen (Krn) from surrounding cells to promote proliferation (Jiang *et al*. 2011). Moreover, suppression of EGFR signaling during homeostasis also leads to a loss of EBs and ISCs (Jiang *et al*. 2011). This suggests that the ECs>ISCs L-R interactions of the EGFR pathway are active during homeostasis, albeit at a lower level. Consistent with this, we observed expression of *Krn* in three EC clusters (aEC4, mEC, and pEC2-3) and *spi* in the ISC/EB cluster. In addition, we also observed *Vn* expression in pEC2 and pEC3. Similar to the JAK/STAT pathway, cell interaction scores for EC to ISC/EB signaling for Vn-, Krn- or Spi-Egfr L-R pairs were greater than those of ISC/EB to EC signaling. This suggests that FlyPhoneDB can accurately predict the directionality of L-R pairs between cell types. Furthermore, EGFR signaling in the midgut is likely to be active at a basal level during homeostasis and could be primed for ISC proliferation in the event of damage.

Interestingly, among the many interactions observed in the FlyPhoneDB output for the midgut dataset, we found very strong interaction scores originating from various EE clusters that were directed towards other cell types. For example, all EE to ISC/EB interaction scores were strong with regards to the Spi-Egfr and Krn-Egfr L-R interactions. This suggests that EEs may send signals to ISCs to promote proliferation. Among other L-R pairs, Tk-TkR99D and Dh31-Dh31R have strong interaction scores between AstA-EE or NPF-EE and many EC or EC-like clusters. The Tk-TkR99D observation is consistent with expectation, as the secretion of TK by EEs is thought to regulate lipid metabolism in ECs (Song *et al*. 2014). Less is known about the interaction of Dh31-Dh31R between EE and ECs, suggesting this is a topic that could be explored further. Finally, we also observed high interaction scores when comparing EEs between itself and other EEs cluster interaction scores for the L-R pair, Sema1a-PlexA. In the nervous system, Sema1a binds to its receptor PlexA and triggers a cascade of downstream events that lead to a repulsive growth cone response (Winberg *et al*. 1998; Winberg *et al*. 2001). This raises the interesting possibility that a similar mechanism is used by EEs to ensure that they are evenly distributed along the midgut. Altogether, these findings show that FlyPhoneDB can generate new predictions that can be further tested experimentally.

To further evaluate the strength of FlyPhoneDB predictions, we next applied the FlyPhoneDB analysis approach to a second dataset. The dataset was generated using single nuclei RNA-seq (snRNAseq) on whole abdomens and predominantly identified three clusters consisting of adipocytes, muscle cells, and oenocytes (Ghosh *et al*. 2020; Ghosh *et al*. 2021). In that study, we reported that the muscle produces the VEGF ligand Pvf1, which acts on its receptor, Pvr, in the oenocytes to inhibit lipid synthesis (Ghosh *et al*. 2020). FlyPhoneDB expression profile analysis accurately identified this cell-cell communication pathway, as Pvf1 and Pvr are enriched in the muscles and oenocytes, respectively. Accordingly, the Pvf1-Pvr interaction score for muscle to oenocyte signaling is greater than found for oenocyte to muscle signaling (Supplementary figure 2).

Finally, we applied FlyPhoneDB to a third study, which surveyed hemocytes in unwounded, wounded, and parasitic wasp-infested larvae (Tattikota *et al*. 2020). The study identified 17 clusters including 12 plasmatocyte (PM) clusters, 2 clusters of lamellocytes (LM), 2 clusters of crystal cells (CC) and one non-hemocyte cluster. Pathway enrichment analysis revealed that the FGF receptor *breathless* (*btl*) and its only ligand, *branchless* (*bnl*), was enriched in the LM2 and CC2 clusters, respectively. Functional analyses further showed that Bnl-Btl L-R communication between LM2-CC2 is crucial for the melanization of parasitoid wasp eggs. Consistent with this, the FlyPhoneDB expression profile indicates that Bnl is enriched in LM2 and Btl in CC2. Furthermore, the Bnl-Btl interaction score for LM2 to CC2 signaling is greater than that of CC2 to LM2 signaling (Supplementary figure 3).

In summary, by testing FlyPhoneDB with existing scRNA-seq datasets for well-studied tissues, we found that we were able to identify known L-R interactions, validating the effectiveness of the approach. In all the cases that we examined, the directionality of L-R between cell-types, when known, was accurately predicted by selecting the higher of the two directional cell-to-cell interaction scores for a pair of cell types. Altogether, our reanalysis using FlyPhone of established directional and autonomous L-R signaling events suggest that the approach can be used to uncover novel cell communication events.

## Discussion

We developed FlyPhoneDB to explore cell-cell communication in *Drosophila* using scRNA-seq data. To provide a high-confidence L-R interaction set, we manually curated L-R interactions from the curated resources, literature and the *Drosophila* community. The strength of FlyPhoneDB analyses depends on an up-to-date database of L-R interactions. Thus, in addition to supporting user-initiated submissions, we will work with FlyBase to curate L-R interactions reported in the literature and import them into FlyPhoneDB. To make FlyPhoneDB user-friendly, we also developed the web-based FlyPhoneDB Explorer, which allows users to easily search the L-R database, upload new L-R annotations, browse the midgut cell crosstalk example, and upload scRNA-seq data to perform their own analyses. We also note that current scRNA-seq data do not provide spatial information and note that FlyPhoneDB can help predict spatial locations of cells in specific clusters by projecting cell-cell communication pathways onto the anatomy.

We made a concerted effort to evaluate all available resources for building FlyPhoneDB, including by importing information from FlyBase, the go-to community resource for annotated information about *Drosophila*. Nevertheless, there remains room for improvement, in particular as new knowledge emerges. To facilitate community updates, we implemented a page at which users can use a form to suggest new L-R pairs and provide the relevant publication(s) supporting the addition. With community input as well as potential annotation pipeline from FlyBase, we anticipate that FlyPhoneDB will improve and expand over time, further increasing its value to the community.

FlyPhoneDB present a number of novel features compared to other analysis tools that predict cell-cell communication from scRNA-seq data. First, we manually curated the data to ensure the accuracy of the L-R database. Second, none of the existing tools currently provide information for core components. Third, FlyPhoneDB displays the activity of core components for each pathway through a heatmap, which can help users quickly compare the pathway activities between different cell types. Finally, we developed the FlyPhoneDB web server that enables users without programming experience to analyze and obtain results through web pages quickly.

## Acknowledgements

We would like to thank the members of the Perrimon laboratory, Drosophila RNAi Screening Center (DRSC), Transgenic RNAi Project (TRiP) for helpful suggestions regarding the project, and Stephanie Mohr for comments on the manuscript. Relevant grant support includes NIH NIGMS P41 GM132087 and BBSRC-NSF. J.S.S.L is supported by a Croucher fellowship for Postdoctoral Research from the Croucher Foundation. H.A. is supported by the British Medical Research Council grant #MR/N030117/1, N.P. is an investigator of Howard Hughes Medical Institute.

---

**Supplementary figure 1.**
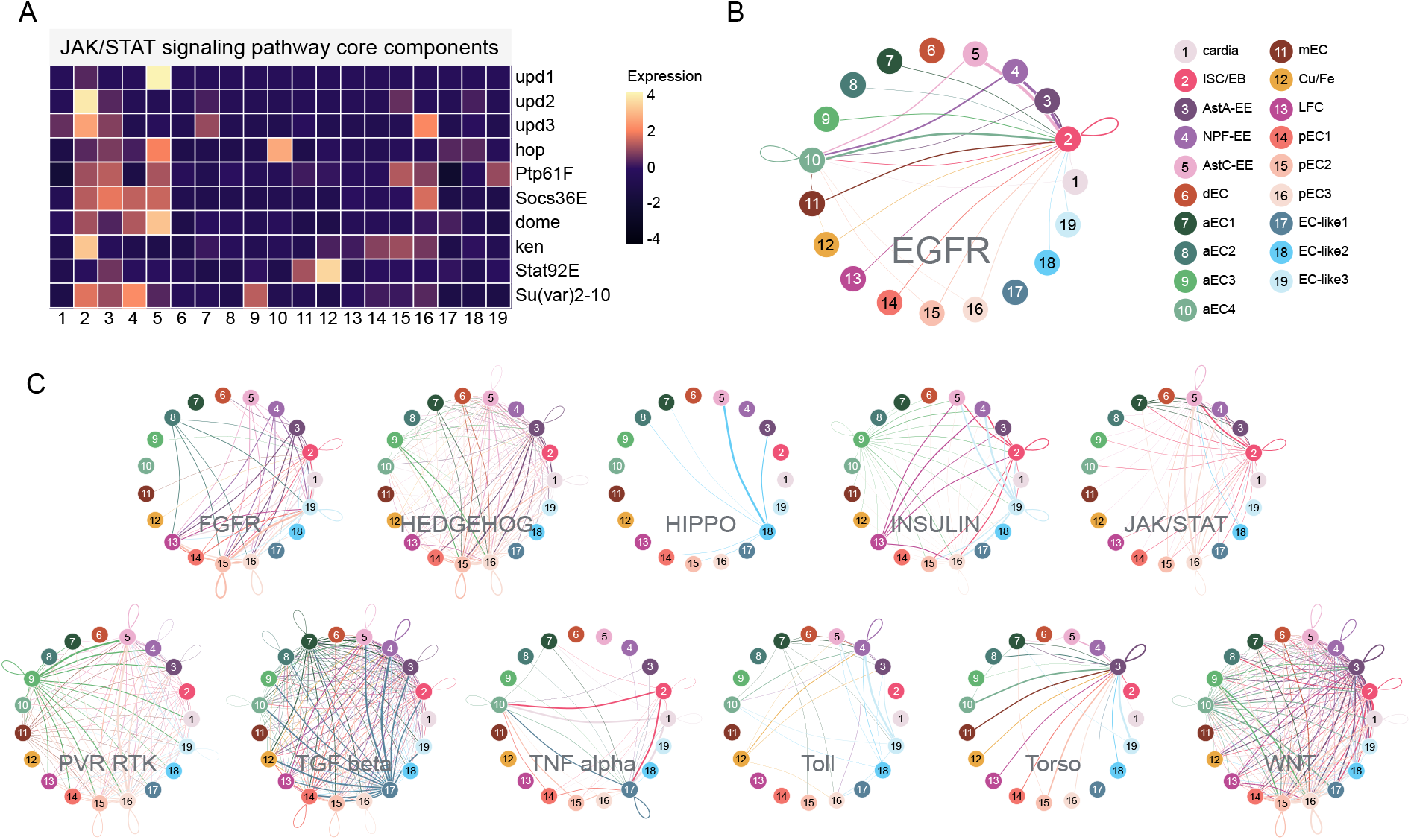
Inferred gut signaling networks. (A) Heatmap shows JAK/STAT signaling pathway core components activity from the Hung *et al*. 2020 dataset (B) Circle plot shows the cell-cell communication for EGFR signaling pathways, providing a summary of cell interactions at the signaling pathway level. (C) Circle plot of the other 11 signaling pathways.

**Supplementary figure 2.**
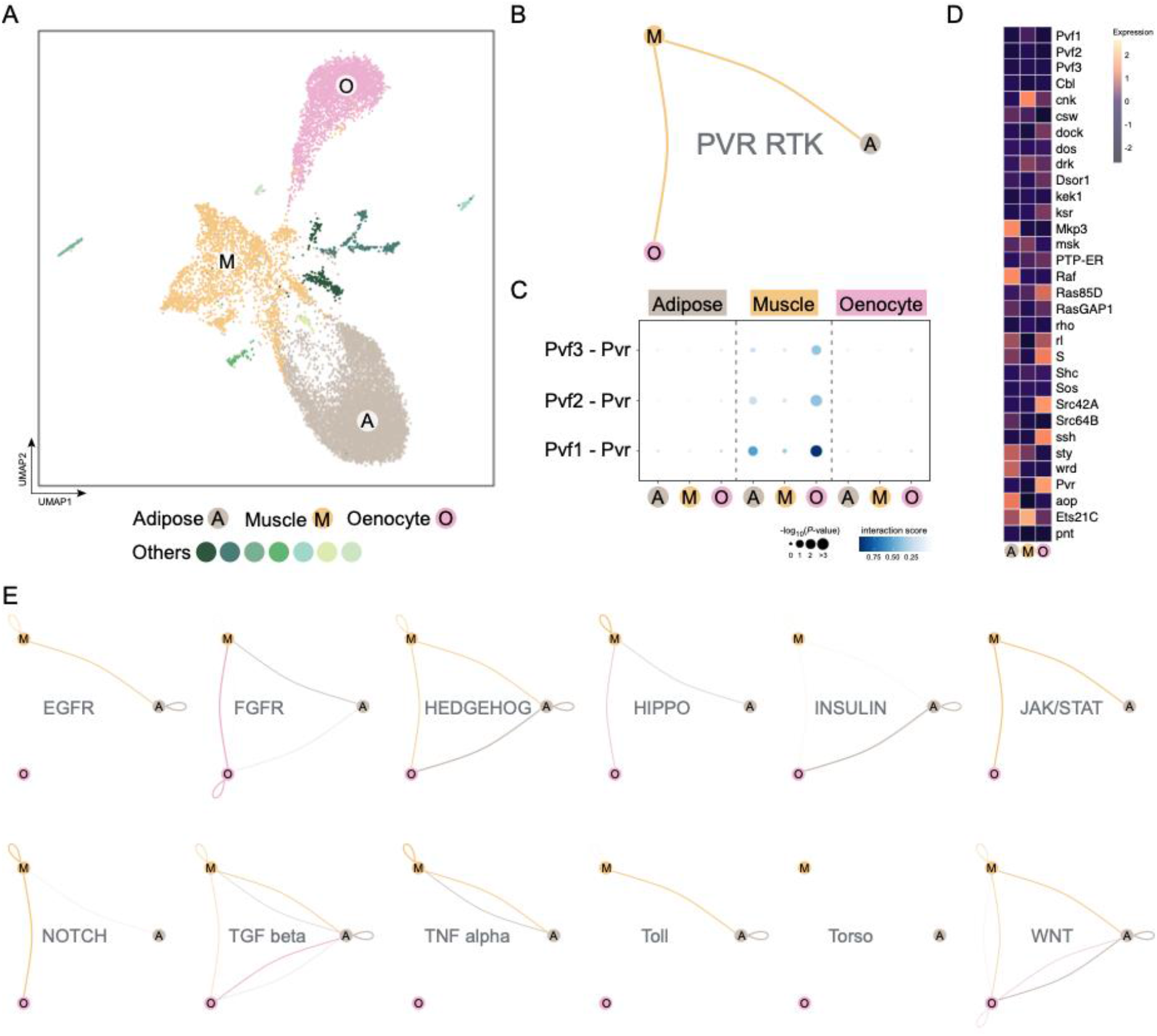
Inferred abdomen signaling networks. (A) UMAP of the *Drosophila* abdomen from Ghosh *et al*. 2020. (B) Circle plot showing the cell-cell communication for PVR RTK signaling pathway. (C) Dot plot showing the strong Pvf1-Pvr interaction from muscle to oenocytes. (D) Heatmap showing PVR RTK signaling pathway core components activity in the abdomen. (E) Circle plot of the other 12 signaling pathways for the Ghosh *et al*. 2020 dataset.

**Supplementary figure 3.**
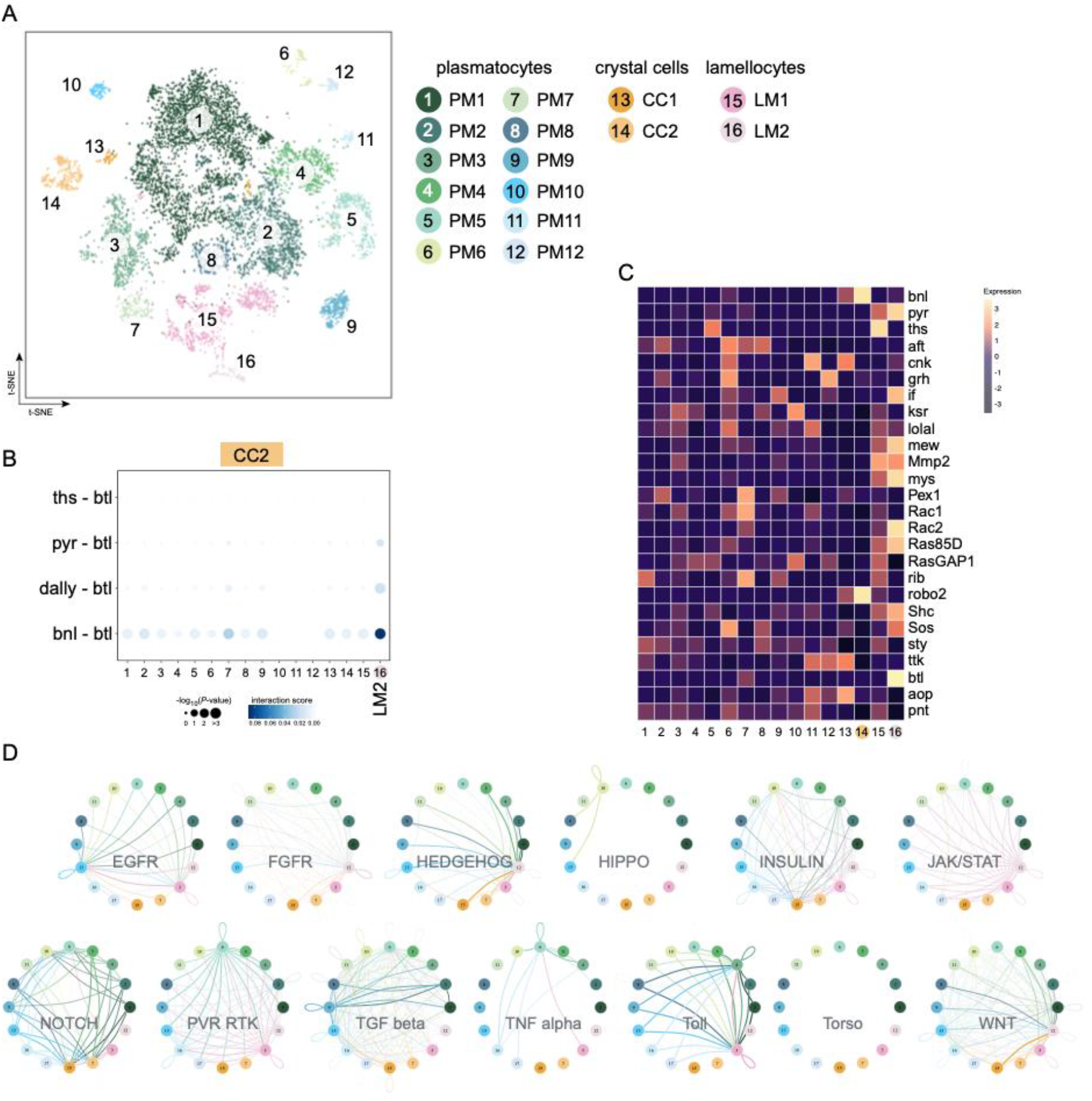
Inferred blood signaling networks. (A) t-SNE of the *Drosophila* blood from Tattikota *et al*. 2020. (B) DotPlot showing the Bnl-Btl interaction from CC2 to LM2. (C) Heatmap showing FGFR signaling pathway core components activity in the blood. (D) Circle plot of the 13 signaling pathways for the Tattikota *et al*. 2020 dataset.

## Literature Cited

Browaeys, R., W. Saelens and Y. Saeys, 2020 NicheNet: modeling intercellular communication by linking ligands to target genes. Nat Methods 17: 159–162.

Buchon, N., D. Osman, F. P. David, H. Y. Fang, J. P. Boquete et al., 2013 Morphological and molecular characterization of adult midgut compartmentalization in Drosophila. Cell Rep 3: 1725–1738.

Dutta, D., A. J. Dobson, P. L. Houtz, C. Glasser, J. Revah et al., 2015 Regional Cell-Specific Transcriptome Mapping Reveals Regulatory Complexity in the Adult Drosophila Midgut. Cell Rep 12: 346–358.

Efremova, M., M. Vento-Tormo, S. A. Teichmann and R. Vento-Tormo, 2020 CellPhoneDB: inferring cell-cell communication from combined expression of multi-subunit ligand-receptor complexes. Nat Protoc 15: 1484–1506.

Fan, H. C., G. K. Fu and S. P. Fodor, 2015 Expression profiling. Combinatorial labeling of single cells for gene expression cytometry. Science 347: 1258367.

Ghosh, A. C., S. G. Tattikota, Y. Liu, A. Comjean, Y. Hu et al., 2020 Drosophila PDGF/VEGF signaling from muscles to hepatocyte-like cells protects against obesity. Elife 9.

Ghosh, A. C., S. G. Tattikota, Y. Liu, A. Comjean, Y. Hu et al., 2021 Correction: Drosophila PDGF/VEGF signaling from muscles to hepatocyte-like cells protects against obesity. Elife 10.

Gontijo, A. M., and A. Garelli, 2018 The biology and evolution of the Dilp8-Lgr3 pathway: A relaxin-like pathway coupling tissue growth and developmental timing control. Mech Dev 154: 44–50.

Guo, X., C. Yin, F. Yang, Y. Zhang, H. Huang et al., 2019 The Cellular Diversity and Transcription Factor Code of Drosophila Enteroendocrine Cells. Cell Rep 29:4172–4185 e4175.

Hu, Y., A. Comjean, L. A. Perkins, N. Perrimon and S. E. Mohr, 2015 GLAD: an Online Database of Gene List Annotation for Drosophila. J Genomics 3: 75–81.

Hu, Y., A. Vinayagam, A. Nand, A. Comjean, V. Chung et al., 2018 Molecular Interaction Search Tool (MIST): an integrated resource for mining gene and protein interaction data. Nucleic Acids Res 46: D567–D574.

Hung, R. J., Y. Hu, R. Kirchner, Y. Liu, C. Xu et al., 2020 A cell atlas of the adult Drosophila midgut. Proc Natl Acad Sci U S A 117: 1514–1523.

Hung, R. J., J. S. S. Li, Y. Liu and N. Perrimon, 2021 Defining cell types and lineage in the Drosophila midgut using single cell transcriptomics. Curr Opin Insect Sci 47: 12–17.

Huntley, R. P., T. Sawford, P. Mutowo-Meullenet, A. Shypitsyna, C. Bonilla et al., 2015 The GOA database: gene Ontology annotation updates for 2015. Nucleic Acids Res 43: D1057–1063.

Jiang, H., M. O. Grenley, M. J. Bravo, R. Z. Blumhagen and B. A. Edgar, 2011 EGFR/Ras/MAPK signaling mediates adult midgut epithelial homeostasis and regeneration in Drosophila. Cell Stem Cell 8: 84–95.

Jiang, H., P. H. Patel, A. Kohlmaier, M. O. Grenley, D. G. McEwen et al., 2009 Cytokine/Jak/Stat signaling mediates regeneration and homeostasis in the Drosophila midgut. Cell 137: 1343–1355.

Jin, S., C. F. Guerrero-Juarez, L. Zhang, I. Chang, R. Ramos et al., 2021 Inference and analysis of cell-cell communication using CellChat. Nat Commun 12: 1088.

Klein, A. M., L. Mazutis, I. Akartuna, N. Tallapragada, A. Veres et al., 2015 Droplet barcoding for single-cell transcriptomics applied to embryonic stem cells. Cell 161: 1187–1201.

Larkin, A., S. J. Marygold, G. Antonazzo, H. Attrill, G. Dos Santos et al., 2021 FlyBase: updates to the Drosophila melanogaster knowledge base. Nucleic Acids Res 49: D899–D907.

Lieber, T., S. Kidd and M. W. Young, 2002 kuzbanian-mediated cleavage of Drosophila Notch. Genes Dev 16: 209–221.

Macosko, E. Z., A. Basu, R. Satija, J. Nemesh, K. Shekhar et al., 2015 Highly Parallel Genome-wide Expression Profiling of Individual Cells Using Nanoliter Droplets. Cell 161: 1202–1214.

Micchelli, C. A., and N. Perrimon, 2006 Evidence that stem cells reside in the adult Drosophila midgut epithelium. Nature 439: 475–479.

Ohlstein, B., and A. Spradling, 2006 The adult Drosophila posterior midgut is maintained by pluripotent stem cells. Nature 439: 470–474.

Osman, D., N. Buchon, S. Chakrabarti, Y. T. Huang, W. C. Su et al., 2012 Autocrine and paracrine unpaired signaling regulate intestinal stem cell maintenance and division. J Cell Sci 125: 5944–5949.

Song, W., J. A. Veenstra and N. Perrimon, 2014 Control of lipid metabolism by tachykinin in Drosophila. Cell Rep 9: 40–47.

Stuart, T., A. Butler, P. Hoffman, C. Hafemeister, E. Papalexi et al., 2019 Comprehensive Integration of Single-Cell Data. Cell 177:1888–1902 e1821.

Tattikota, S. G., B. Cho, Y. Liu, Y. Hu, V. Barrera et al., 2020 A single-cell survey of Drosophila blood. Elife 9.

Upadhyay, A., L. Moss-Taylor, M. J. Kim, A. C. Ghosh and M. B. O’Connor, 2017 TGF-beta Family Signaling in Drosophila. Cold Spring Harb Perspect Biol 9.

Winberg, M. L., J. N. Noordermeer, L. Tamagnone, P. M. Comoglio, M. K. Spriggs et al., 1998 Plexin A is a neuronal semaphorin receptor that controls axon guidance. Cell 95: 903–916.

Winberg, M. L., L. Tamagnone, J. Bai, P. M. Comoglio, D. Montell et al., 2001 The transmembrane protein Off-track associates with Plexins and functions downstream of Semaphorin signaling during axon guidance. Neuron 32: 53–62.

